# Valid, invalid, or somewhere in between? Baseline ImPACT and stand-alone performance validity testing in collegiate athletes

**DOI:** 10.1101/2023.05.03.538988

**Authors:** Kate L. Higgins, Heather C. Bouchard, Julia E. Maietta, Julia M. Laing-Young, Douglas H. Schultz

**Author notes:** Correspondence concerning this article should be addressed to: Kate Higgins, UNL Athletics, 800 Stadium Drive, Lincoln, NE 68588, United States. Author Note The authors have no conflicts of interest to disclose.

## Abstract

**Objective:** Baseline cognitive testing is important for sport concussion management. Assessing baseline data for both the validity and reliability is vital to ensuring its utility. Stand-alone performance validity tests (PVT) and embedded validity indicators (EVI) (collectively called “validity measures” hereafter) are commonly used in neuropsychological testing and screening. This study further investigates efficient ways to assess validity of baseline cognitive testing.

**Methods:** 231 NCAA athletes completed baseline assessment with ImPACT and one of two PVT’s: Medical Symptom Validity Test or Rey Dot Counting Test. The relationship between scores on validity measures and cognitive performance, symptoms, and sleep was assessed.

**Results:** Athletes who failed two or more validity measures performed worse on the Visual Motor Speed Composite while athletes failing three or more measures performed worse on the Reaction Time Composite. Those athletes who failed validity measures and also reported poor sleep performed worse on both composites. Self-reported symptoms and hours of sleep were not independently associated with failing validity measures. Lastly, athletes with self-reported ADHD and/or LD were more likely to fail two+ validity measures (46% versus 27% of neurotypical athletes).

**Conclusions:** Consistent with previous research, ImPACT Baseline++ only identified 1.7% of the sample’s data as invalid despite 4.8% of the dataset failing a combination of PVT and EVI and an additional 42.9% failing at least EVI alone. Results suggest that assessing validity on a continuum and using multiple validity measures may be useful to identify data validity that falls in the middle of the continuum.

**Public Significance Statement:** Baseline concussion testing is common and often mandated prior to sport participation, however, a baseline only has utility if it is both a reliable and valid representation of the athlete’s healthy and optimal functioning. This study adds to the growing body of literature demonstrating that baseline testing may frequently have questionable validity. It also provides support to the value of conceptualizing validity as a spectrum, rather than dichotomy and is the first to apply the concept to baseline concussion testing data.

---

Concussion baseline testing with some combination of cognitive, symptoms, and balance/vestibular/oculomotor evaluation has become common and is required at many levels of sport participation. At the NCAA level, the evaluation of these three elements are required for varsity collegiate athletes as a part of their pre-participation evaluation (National Collegiate Athletic Association Sport Science Institute, 2022).

The premise of baseline testing is that an athlete can serve as their own standard for “recovered” should they receive a concussion (Bailey et al., 2009). However, variability in the testing environment or in individual factors, including effort, can impact the validity of baseline testing scores resulting in a potentially inaccurate standard for “normalized” functioning after a concussion. While there is mixed evidence regarding the rate of invalid performance in athletes on baseline cognitive testing, a recent systematic review (Gaudet & Weyandt, 2017) found that between 2.7% and 27.9% have indications of invalidity using a variety of validity indicators within Immediate Post-Concussion Assessment and Cognitive Testing (ImPACT) a computerized neurocognitive screener commonly used for baseline concussion evaluations in the sport setting (Gaudet & Weyandt, 2017). Invalid baseline has serious clinical implications. If an athlete’s data does not provide an accurate representation of their individual baseline functioning, they are at risk of being returned to sport and head injury exposure before they are fully recovered, which puts them at increased risk for a subsequent concussion (Bailey et al., 2006). As a result, assessing for validity in the context of baseline testing has begun to be addressed in the literature, particularly around ImPACT. ImPACT contains several “built-in” embedded performance validity indicators, with associated cut-scores, that are used by the program to identity invalid baselines (labeled “Baseline++”). As will be discussed further, these do not appear to be particularly sensitive to invalid performance, missing up to 20% of test-takers who are instructed to intentionally underperform on ImPACT testing (Gaudet & Weyandt, 2017). Because of this, additional embedded validity indicators have been identified in the literature to improve the sensitivity of invalid baseline classification (Higgins et al., 2017; Manderino & Gunstad, 2018; Manderino et al., 2019; Schatz & Glatts, 2013), but have not been incorporated into the ImPACT program.

Assessing for performance validity is a common and important aspect of traditional, paper-and-pencil neuropsychological assessment and there are a multitude of factors that can contribute to less-than-optimal performance. These factors include both individual factors (including effort provided by the examinee, lack of sleep, illness- Bailey et al., 2006) and environmental factors (including distractions and malfunctioning technology- Moser et al., 2015). Validity testing is often colloquially, and inaccurately, called “effort testing”. The inaccuracy in this label is that performance validity testing cannot identify the cause of the invalidity in the data, just its presence or lack thereof. While the cause of invalid data is certainly clinically relevant in individual patient care, identifying the factors contributing to invalidity in a single dataset must be left up to the expertise of the individual clinician, as it is beyond the capacity of performance validity testing. Given that the myriad of factors that can affect data validity may change between individuals and testing sessions, or even within an individual within a testing session, determining the cause or intention of invalid performance on a group level (such as assuming that all invalid baseline testing is due solely to a lack of effort) is beyond the scope of performance validity testing

Valid performance is characterized by full engagement and best performance by the examinee and no interference of any kind from the testing environment. At the other end of the validity spectrum is malingering (i.e., intentional poor effort for some external incentive), or in sport concussion contexts, sandbagging (intentionally performing poorly to be cleared for return-to-play status more quickly after concussion). Between the two poles of this validity spectrum is a large space where performance is suboptimal, so data is not valid, but is conceptually different that intentional underperformance (Erdodi, 2023; Heilbronner et al., 2009; Sweet et al., 2021).

In traditional neuropsychological evaluations, it is common to employ a combination of stand-alone performance validity tests (PVT)- tests conducted for the sole purpose of evaluating performance validity- and embedded performance validity indicators (EVI)- scores from a test given to measure a different cognitive domain (Heilbronner et al., 2009; Miller et al., 2017; Sweet, Heilbronner, et al., 2021). While both types of validity measures are often employed in neuropsychological testing, there is a breadth of literature demonstrating higher sensitivity and specificity of single stand-alone PVTs for performance invalidity compared to EVIs and use of PVT is considered ideal standard practice in recent consensus documents (Curtis et al., 2008; Kanser et al., 2019; Ovsiew et al., 2020; Pliskin et al., 2021; Soble et al., 2021; Sweet, Heilbronner, et al., 2021; Whiteside et al., 2015; Wolfe et al., 2010). But as demonstrated by Erdodi (2023), combining multiple EVI’s can approach both the sensitivity and specificity of stand-alone PVTs and adding a PVT in baseline concussion testing settings is often unrealistic. A survey of health care professionals that manage sport-related concussions revealed that it is not common to use either stand-alone PVTs or thorough evaluation of additional EVIs with ImPACT (Covassin et al., 2009). Gaudet & Weyandt (2017) found, in a systematic review of the literature, that ImPACT Baseline++ EVIs may miss up to 20% of “fake bad” simulators. Taken in conjunction with a general lack of external PVT assessment use, this data suggests some athletes may have baselines that do not reflect their optimal functioning. More recent literature suggests that between 50% and 60% of simulators remain undetected by ImPACT Baseline++ implying a need to re-evaluate the built-in ImPACT EVIs (Abeare et al., 2019; Messa et al., 2020; Raab et al., 2020). Other literature has identified additional embedded indicators of invalidity within ImPACT with greater utility than those used by Baseline++. Specifically, Schatz & Glatts (2013) found that the total number of correct distractors for both Word Memory (< 22) and Design Memory (< 16) identified 90 to 100% of simulated sandbagging. Additionally, Higgins et al. (2017) identified a logistic regression equation that demonstrated 100% sensitivity to simulated sandbagging.

Several studies address the use of stand-alone PVTs in addition to ImPACT testing (Abeare et al., 2019; DaCosta et al., 2021; L. Manderino & Gunstad, 2018; Salazar et al., 2017; Schatz & Glatts, 2013). Specifically, Schatz & Glatts (2013) found that the Medical Symptom Validity Test (MSVT) outperformed the built-in ImPACT EVIs in an experimental sample of college-aged naive and coached malingerers/sandbaggers. The MSVT identified 80 to 90% of naive and coached malingerers compared to 60 to 75% identified by the ImPACT EVIs, respectively. Manderino & Gunstad (2018) also employed an experimental malingering/sandbagging paradigm and found that built-in ImPACT EVIs demonstrated much lower sensitivity than a stand-alone PVT (Word Memory Test) (i.e., the average sensitivity of ImPACT EVIs was 0.25 compared to 0.52 for WMT). Two studies have investigated the use of the Dot Counting Test (DCT) in sport concussion baseline testing (DaCosta et al., 2021; Salazar et al., 2017). Salazar et al. (2017) used naive and coached malingering/sandbagging groups to establish the DCT as an acceptable measure of performance invalidity in this population. DaCosta et al. (2021) used a naturalistic sample of collegiate athletes’ baselines and found that the DCT was more sensitive in detecting invalid cases than built-in ImPACT EVIs alone. Their sample also included a small subsample of athletes with self-reported attention-deficit/hyperactivity disorder (ADHD) or learning disorders (LD), who were more likely to be flagged by ImPACT EVIs but did not fall below the validity cut score on the DCT. DaCosta et al. (2021) is one of the first to compare performance on ImPACT EVIs to stand-alone PVTs in a population with neurodevelopmental diagnoses. There is additional research which has similarly suggested that athletes with neurodevelopmental disorders may be flagged as invalid on ImPACT more frequently than their neurotypical counterparts (Gaudet & Weyandt, 2017; Maietta, Barchard, et al., 2021; Maietta et al., 2023). Theories to explain these findings include athletes with self-reported ADHD and/or LD may fall below EVI thresholds due to underlying reading and/or attentional difficulties rather than intentional attempts to underperform.

Consistent with DaCosta et al. (2021)’s findings, there is additional evidence that individuals with confirmed clinical diagnoses of ADHD and/or LD may not fail other stand-alone and embedded validity indicators at higher rates, but rather have lower rates of PVT failure than other clinical populations (Pearson, 2009). These findings suggest that the higher rates of validity failure in athletes with self-reported ADHD and/or LD may be specific to ImPACT’s EVIs. Further research is needed in this regard given these findings and the importance of understanding PVT performance in these populations.

The focus on experimental malingering/sandbagging paradigms in all but one study in the extant literature may be problematic, as these patterns of performance may produce exaggerated profiles that are easily detected by traditional PVTs. Both sandbagging and malingering involve the intention to underperform. This intentionality may not be present in all athletes, and may be better explained by the concept of effort existing on a continuum (in contrast to as a dichotomy) initially laid out in the American Academy of Clinical Neuropsychology’s (AACN) 2009 consensus statement on validity assessment (Heilbronner et al., 2009) and further discussed in their 2021 follow-up consensus statement (Sweet, Heilbronner, et al., 2021). There is also evidence that many athletes may underperform with more subtle methods than those used by research participants simulating sandbagging or by patients actively attempting to malinger during neuropsychological testing (Abeare et al., 2019). Indifference, dislike of the test due to multiple exposures, or lack of understanding of ImPACT baseline’s utility in the management of sport concussion may also play a role in an athlete’s engagement in testing or variability in engagement across the course of testing (Erdal, 2012; McClure et al., 2014; Rabinowitz et al., 2015; Schatz et al., 2017; Walton et al., 2018). If the effort supplied by an athlete falls on the middle of the spectrum (rather than on the “good” end or the “poor” end), there may also be more interference in cognitive scores from poor sleep, emotional distress, cognitive fatigue, etc. given there is less engagement to help mitigate the effects of these factors. The concept of “transient validity” is also raised in the AACN 2021 follow-up consensus statement (Sweet, Heilbronner, et al., 2021). This describes failure on a single validity measure in the absence of intentional underperformance. This concern supports the use of multiple validity measures (both embedded and stand-alone) throughout a neuropsychological evaluation. While baseline concussion testing is significantly shorter than a full neuropsychological evaluation, the concept of multiple validity checks may be useful in differentiating less intentional “middle of the validity spectrum” performance from intentional underperforming. Neurodevelopmental diagnoses, such as ADHD or LD, can make the classification of validity even more variable, particularly when using ImPACT EVIs (DaCosta et al., 2021; Maietta et al., 2021; Maietta et al., 2023; Manderino et al., 2019). These factors individually or in combination could easily play a role in athletes providing less-than-valid data without intentionally underperforming or sandbagging.

Therefore, there is a need to further investigate efficient ways to evaluate performance validity in baseline cognitive concussion assessment through either stand-alone PVTs, EVIs in ImPACT, or a combination of the two. Given the well documented utility of stand-alone PVTs in detecting intentional underperformance in traditional neuropsychological batteries, adding a stand-alone PVT to supplement baseline ImPACT in the detection of invalid performance is consistent with best practice guidelines in the field (Abeare et al., 2019; DaCosta et al., 2021; Sweet, Heilbronner, et al., 2021), but questions remain about their utility in detecting less-than- valid data that may be related to factors other than intentional underperformance and the return-on-investment of adding an additional measure to concussion baselines. Therefore, this research will evaluate the rates of failed effort measures in a baseline concussion testing sample, as well as the agreement (or lack of agreement) between embedded ImPACT EVIs and stand-alone “gold-standard” PVT measures in identifying invalidity to determine the utility of adding a PVT to baseline testing.

## Methods

### Participants

NCAA Division I collegiate athletes completed baseline assessments annually (with two years of data examined for this research) as part of their standard clinical pre-participation evaluations. All athletes completed ImPACT (*N* = 231). In addition to ImPACT, athletes completed the Medical Symptom Validity Test (*n* = 130) in year one, and athletes completed the Rey Dot Counting Test (*n* = 101) in year two. Ages ranged from 17 to 24 (*M =* 18.6*, SD =* 1.3; 63% male). Race and ethnicity were self-reported by participants, and at the time of data collection, were collected as a single variable instead of two. Participants self-identified as White (67%), African American/Black (17%), Biracial/Multiracial (10%), Latinx/Hispanic (4%), and Asian (2%). The majority of participants were native to the United States (92%). A subset of participants self-reported diagnoses of ADHD, LD, or a combination of the two - hereafter designated self-reported ADHD and/or LD (*n* = 44). Participants were from 15 sports, including baseball/softball (n = 27), basketball (n = 17), bowling (n = 3), cross-country (n = 13), football (n = 68), golf (n = 10), gymnastics (n = 14), rifle (n =4), soccer (n = 9), swimming (n = 10), tennis (n = 6), track & field (n = 36), volleyball (n = 3), and wrestling (n = 11).

### Materials

#### ImPACT

ImPACT is a computerized neuropsychological assessment used to assess cognitive performance at baseline and recovery following concussion by comparing post-concussion scores to baseline performance (ImPACT Applications, 2021). Four composite scores (Verbal Memory, Visual Memory, Visual-Motor Speed, and Reaction Time) are calculated from 10 cognitive tasks. In addition to cognitive assessment, athletes self-report their total symptoms, and optionally report their total hours of sleep the night prior to testing. EVIs built into ImPACT (Baseline++) are: Impulse Control Composite > 30, Word Memory Learning Percent Correct < 69%, Design Memory Learning Percent Correct < 50%, Three Letters Total Letters Correct < 8 (ImPACT Applications, 2021) and X’s and O’s Total Incorrect > 30 (ImPACT Applications, 2016). Additional EVIs (that are not built into ImPACT) include: Word Memory Correct Distractors (WMCD) < 22 and Design Memory Correct Distractors (DMCD) < 16 (Schatz & Glatts, 2013) and a logistic regression equation (LRE) using Word Memory Learning Percent Correct, Word Memory Delay Memory Percent Correct, Design Memory Total Percent Correct, and X’s and O’s Total Correct (Interference) (Higgins et al., 2017). When evaluating cognitive tasks, only the full composites for Visual Motor Speed and Reaction Time were evaluated, as portions of both Verbal Memory and Visual Memory composites are used in the EVIs described above. Individual variables from each of Verbal Memory and Visual Memory composites were evaluated as a proxy (Three Letters-Total Sequence Correct for Verbal Memory composite and X’s and O’s- Total Correct/Memory for Visual Memory composite) as these variables are part of the composites, but are not included in any of the EVIs assessed in the following analysis.

#### Rey Dot Counting Test (DCT)

The Rey Dot Counting Test (DCT) is a timed task designed by Rey (1941) to detect performance invalidity with a total administration time of less than 5 minutes. The test consists of twelve index cards containing differing numbers of dots. Several scores are calculated from the test: mean counting times, ratio (mean time to count different groups of dots), errors (total number of errors), and a combination score (mean dot counting time for each group + number of errors) (Boone et al., 2002). A cut-score of ≥15 was used to identify performance invalidity, based on previous research with a collegiate athlete population (DaCosta et al., 2021).

#### Medical Symptom Validity Test (MSVT)

The MSVT (Green, 2004) is a computerized, stand-alone PVT. The MSVT takes approximately 15-20 minutes to complete, including a 10- minute delay and delayed memory trials. A list of related word pairs are presented on a computer screen and then the participant has to identify the learned words using a variety of memory tasks. Immediate Recall, Delayed Recall, and Consistency variables were analyzed with ≤ 85% on any of the three variables signifying performance invalidity (Green, 2004).

### Procedure

As part of their standard clinical pre-participation concussion evaluation, new student-athletes completed a series of brief assessments, including PVTs, administered by either a sports neuropsychologist or trained doctoral students. ImPACT test administration occurred in a group setting using a standardized administration protocol. PVT’s were completed in an individualized setting. ImPACT and the PVT included in baseline assessment for that year (either DCT or MSVT) were all administered on the same day, but possibly not back-to-back (due to administration order of the multiple assessments completed as a part of concussion baseline evaluation). The use of student athletes’ deidentified data for research purposes, and the use of these data for this specific study, was approved by the home institution’s Institutional Review Board (IRB).

### Statistical Analysis

Statistical analyses were conducted in R (RStudio Team, 2021). The relationship between validity measures and clinical outcomes, specifically cognitive task scores, total symptom score, and sleep was assessed using linear models with the *lm* command. Performance validity was evaluated across a continuum (zero, one, two and three+ failed validity measures) within the linear models, except in the interaction models. In these models, validity was evaluated as a binary variable (zero or one+ failed validity measures). Linear models were also used to assess the relationship between self-reported ADHD and/or LD diagnosis and clinical outcomes. The relationship between categorical variables, including performance validity and self-reported ADHD and/or LD diagnosis, was assessed using a chi-square test of independence with the *chisq.test* commands. Effect sizes were calculated for linear regressions using the Claudy-3 formula for models with four predictors (e.g., zero, one, two, or three+ failed measures) or with the Smith & Wherry formula for models with two predictors (e.g., ADHD/LD)(Yin & Fan, 2001). The effect sizes were calculated with the *effect.size* command from the *yhat* package.

## Results

Records for 231 collegiate athletes’ clinical concussion baseline assessments were evaluated. Of the 231 athletes, 110 failed at least one validity measure (either on EVI, PVT, or a combination of the two). Thirty-nine athletes failed only one validity measure, 40 athletes failed two validity measures, and 31 athletes failed three or four (three+) validity measures. Table 1 displays these results and the number of athletes who failed each individual validity measure. Neither self-reported total symptoms (*F*(3, 227) = 0.92, *p* = 0.431) nor self-reported hours of sleep the previous night (*F*(3, 184) = 1.18, *p* = 0.318) were significantly associated with failing one, two, or three+ validity measures.

**Table 1.**
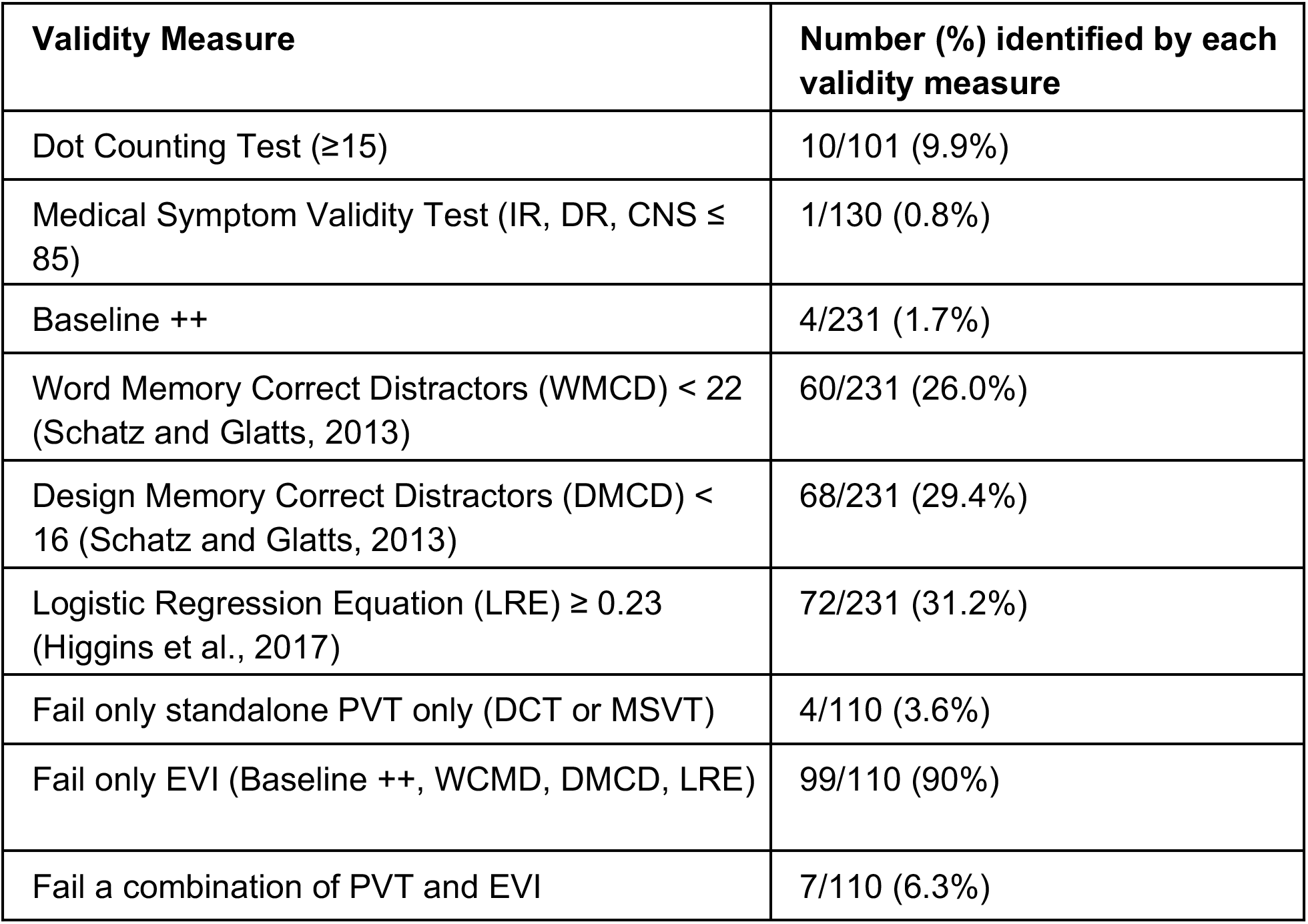
Number of failed effort indicators across individual validity measures.

As expected, the number of failed validity measures was related to cognitive task scores. Visual-Motor Speed composite was associated with failed validity measures (*F*(3, 227) = 9.90, *p* < 0.0001, effect size = 0.10). Figure 1A displays the relationship between Visual-Motor Speed composite performance and number of failed validity measures. Visual-Motor Speed performance was significantly worse for athletes who failed two (*t* = -3.179, *p* = 0.002) or three+ (*t* = -4.866, *p* < 0.0001) validity measures compared to those who did not fail any or for athletes who failed two (*t* = -2.274, *p* = 0.024) or three + (*t* = -3.788, *p* < 0.001) validity measures compared to those who only failed one validity measure. Visual-Motor Speed composite performance was not different for those who failed only one validity measure compared to those who did not fail any (*t* = -0.369, *p* = 0.712) nor was the performance for those who failed two validity measures compared to those who failed three+ (*t* = -1.671, *p* = 0.096). Hours of sleep reported also played a role in Visual-Motor Speed composite scores (*F*(3, 184) = 6.46, *p* < 0.001, effect size = 0.08). Figure 1B displays the interaction between binary validity (i.e., zero failed validity measures and one+ failed validity measures) and self-reported sleep the previous night in relation to Visual-Motor Speed performance. Athletes’ Visual-Motor Speed performance was slower when they both failed validity measures and got less sleep compared to athletes who also reported less sleep, but did not fail any validity measures (*t* = 2.117, *p* = 0.036).

**Figure 1.**
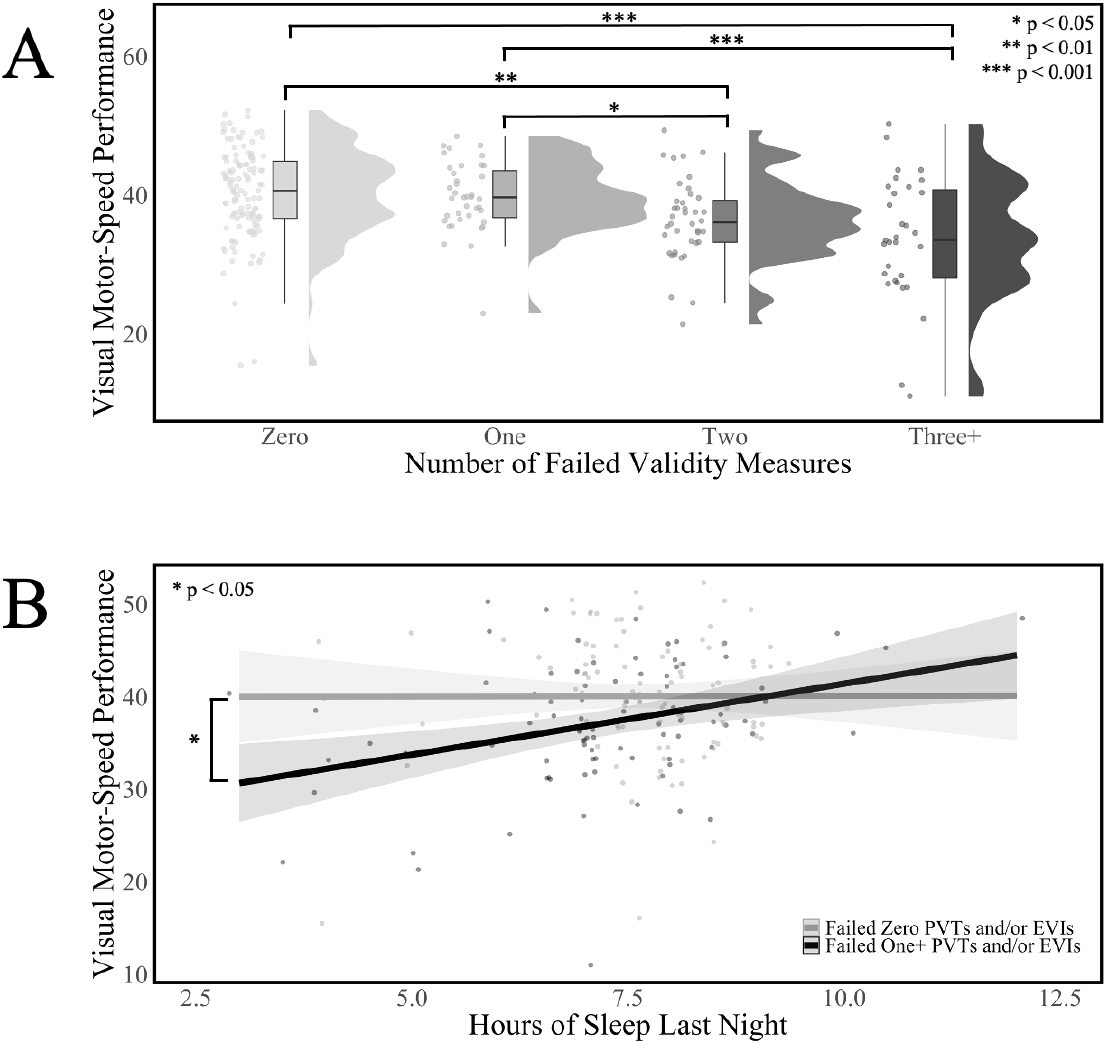

Number of failed validity measures was also associated with Reaction Time composite (*F*(3, 227) = 5.46, *p* = 0.001, effect size = 0.06). Figure 2A displays the relationship between Reaction Time composite performance and number of failed validity measures. Reaction Time composite performance was worse for athletes who failed three+ (*t* = -4.038, *p* < 0.0001) validity measures compared to those who did not fail any validity measures, failed only one validity measure (*t* = -2.621, *p* = 0.009), or failed two validity measures (*t* = -2.527, *p* = 0.012). Additionally, failing one (*t* = 0.990, *p* = 0.323) or two (*t* = 1.141, *p* = 0.255) validity measures did not impact performance on the Reaction Time composite compared to those who did not fail any validity measures. Figure 2B displays the interaction between binary validity and self-reported sleep in relation to Reaction Time performance (*F*(3, 184) = 7.49, *p* < 0.0001, effect size = 0.10). Reaction Time performance was slower in athletes who both reported less sleep and failed one+ validity measure (*t* = -2.730, *p* = 0.007) compared to failing zero validity measures.

**Figure 2.**
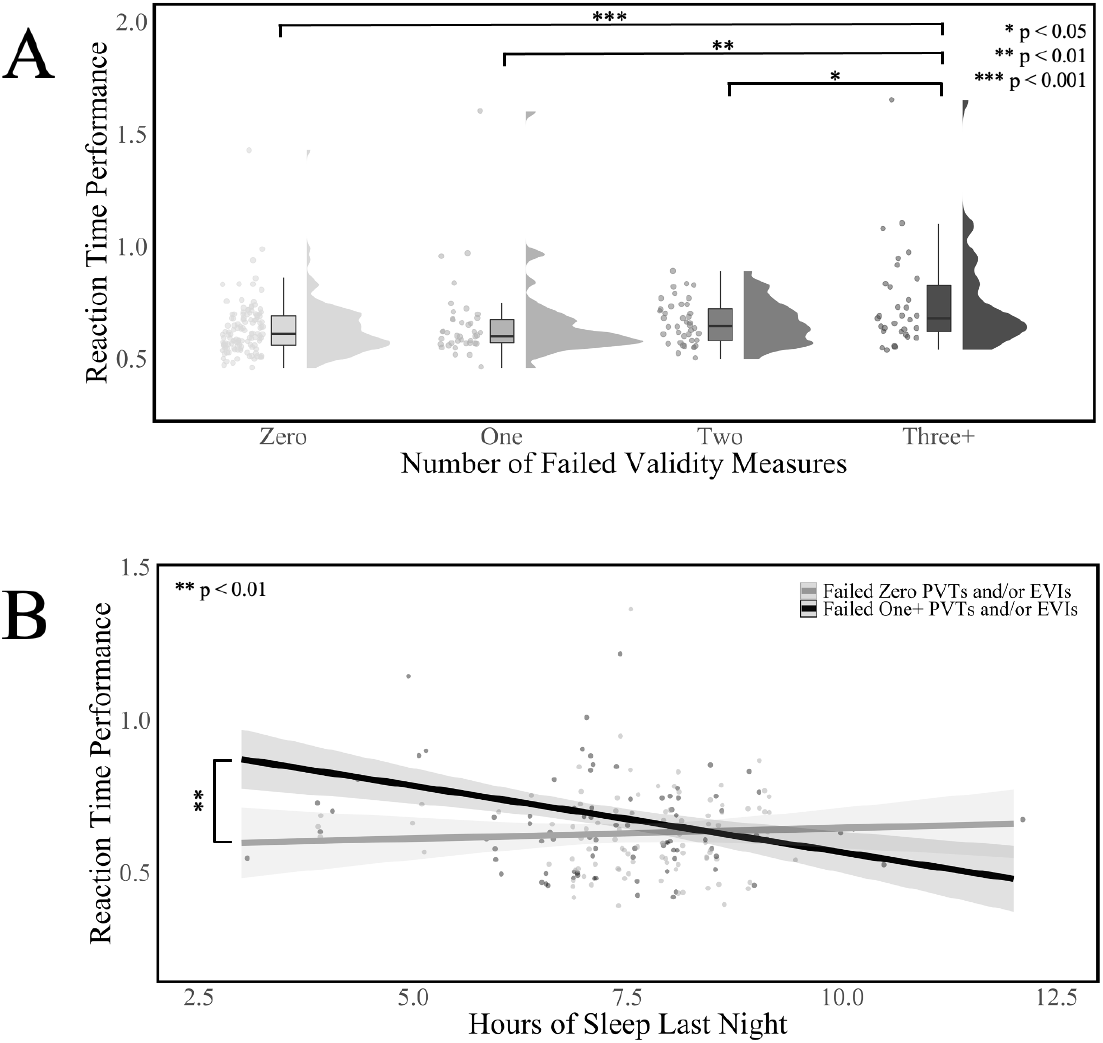

Individual subtest scores- XO Total Correct and Three Letters Total Correct- were used as a proxy for Visual Memory and Verbal Memory composites, respectively, as several of the scores for these composites are used to calculate the embedded PVTs. The number of failed validity measures was significantly associated with the proxies for Verbal Memory (*F*(3, 227) = 4.66, *p* = 0.004, effect size = 0.05) and Visual Memory (*F*(3, 227) = 4.95, *p* = 0.002, effect size = 0.05) performance. Scores on the Verbal Memory composite proxy were significantly worse for athletes who failed two (*t* = -2.483, *p* = 0.014) or three+ (*t* = -2.912, *p* = 0.004) validity measures compared to those who did not fail any. Failing only one validity measure was not significantly different from failing zero validity measures (*t* = -0.335, *p* = 0.752), but it was significantly different from failing two (*t* = -2.271, *p* = 0.024) or three+ (*t* = -2.678, *p* = 0.008) validity measures. Performance on the Visual Memory composite proxy was also significantly worse for athletes who failed two (*t* = -2.253, *p* = 0.025) or three+ (*t* = -3.541, *p* < 0.001) validity measures compared to those who did not fail any. Failing only one validity measure was not significantly different from failing zero (*t* = 0.797, *p* = 0.426) or two validity measures (*t* = - 1.174, *p* = 0.242), it was significantly different from failing three+ (*t* = -2.352, *p* = 0.020).

Self-reported ADHD and/or LD diagnostic status also played a role in ImPACT outcomes. Figure 3 displays the relationship between self-reported ADHD and/or LD diagnosis with self-reported symptoms, self-reported hours of sleep, and Visual-Motor Speed performance. Athletes who self-reported a diagnosis of ADHD and/or LD reported more symptoms (*t* = 4.071, *p* < 0.0001, effect size = 0.06), less sleep (*t* = -2.931, *p* = 0.004, effect size = 0.03), and performed worse on the Visual-Motor Speed composite (*t* = -1.989, *p* = 0.048, effect size = 0.01). However, athletes with a self-reported ADHD and/or LD diagnosis did not perform significantly worse on the Reaction Time composite (*t* = 1.622, *p* = 0.106) compared to athletes who did not endorse a diagnosis.

**Figure 3.**
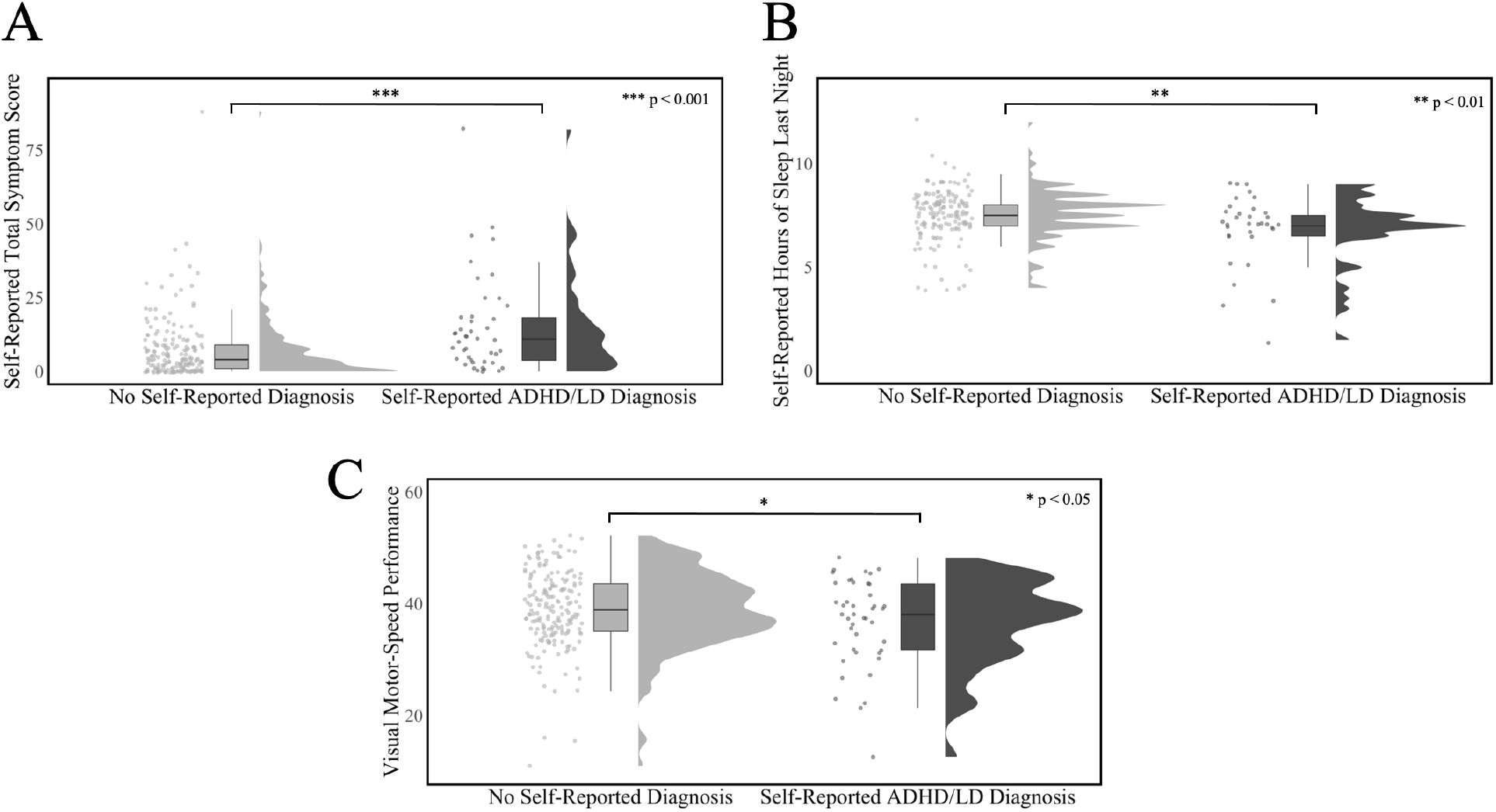

Table 2 reports information regarding athlete validity and self-reported ADHD and/or LD diagnosis. The proportion of athletes who failed one or more validity measures was related to a self-reported diagnosis of ADHD and/or LD (*X^2^* (1, *N* = 231) = 4.83, *p* = 0.028) and athletes who self-reported a diagnosis of ADHD and/or LD were more likely to fail one+ validity measures. A preliminary analysis of athletes with a self-reported diagnosis of ADHD and/or LD (n = 44), assessed validity across a continuum (zero, one, and two+ failed validity measures). The proportion of athletes diagnosed with self-reported ADHD and/or LD significantly differed by validity group (*X^2^*(2, *N* = 231) = 6.55, *p* = 0.038). The proportion of athletes failing only one validity measure was similar (18.2% of athletes with a self-reported diagnosis of ADHD and/or LD vs 16.6% of athletes with no self-reported diagnosis). In contrast, 45.5% of athletes who self-reported a diagnosis of ADHD and/or LD failed two or more validity measures, while only 27.3% of athletes without a self-reported diagnosis of ADHD and/or LD failed two or more validity measures.

**Table 2.** Failed validity measures and self-reported ADHD and/or LD diagnosis - N (%)

| **Number of Failed validity measures** | **No diagnosis** | **Self-reported ADHD and/or LD diagnosis** |
| --- | --- | --- |
| Zero | 105/187<br>(56.1%) | 16/44 (36.4%) |
| One+ | 82/187 (43.9%) | 28/44 (63.6%) |
| **Effort Across a Continuum** |  |  |
| Zero | 105/187<br>(56.1%) | 16/44 (36.4%) |
| One | 31/187 (16.6%) | 8/44 (18.2%) |
| Two | 51/187<br>(27.3%%) | 20/44 (45.5%) |
| **PVTs vs EVIs** |  |  |
| Failed PVT | 7/187 (3.7%) | 4/44 (9.1%) |
| Failed EVI | 79/187 (41.7%) | 27/44 (61.4%) |

## Discussion

Identifying baseline concussion testing data that falls between the “valid” and “intentional underperforming” poles of the validity continuum continues to be challenging, but the implications of data that does not represent an athlete’s optimal functioning is a problem that cannot be ignored. Consistent with previous research (Gaudet & Weyandt, 2017), the embedded Baseline++ validity indicators built into ImPACT appear to do a poor job of identifying invalid data and identified only 1.7% of our sample’s data as invalid despite 4.8% of the sample failing a combination of stand-alone PVT and EVIs and an additional 42.9% failing at least one EVI. In contrast to previous research on concussion baselines and neuropsychological testing in general (Boone et al., 2002; DaCosta et al., 2021; Green, 2004; Schatz & Glatts, 2013), the prevalence of invalid data identified in our sample by stand-alone PVTs was lower than expected. Consistent with DaCosta et al. (2021), there was no overlap between those athletes identified by Baseline++ and PVTs. This lack of overlap may be explained by the very low rates of cases identified as invalid by Baseline++ and stand-alone PVTs- 1.7% and 3.6% of the sample respectively- although this raises the question of the validity of the measures given that they are supposed to be measuring the same construct.

Also consistent with previous research, the EVIs suggested by Higgins et al. (2017) and Schatz & Glatts (2013) identify a notably larger number of athletes’ data as questionably valid than Baseline++ or stand-alone PVTs. The lack of sensitivity in the PVTs to invalidity in the data from this population is not surprising, as collegiate athletes are a relatively cognitively high-functioning population. PVTs are created so that cognitively impaired individuals can pass, so PVTs are not calibrated to catch subtle variations in validity that could be demonstrated by a high functioning sample. While the cut-offs utilized in this research are those suggested by either the test manual (MSVT) or relevant research (DCT), more stringent cut-offs may have increased the sensitivity of the stand-alone PVTs utilized. Ongoing evaluation of the EVI’s identified in the research is also warranted. While they have high face validity in that they were developed through instructing the research participants- and in some cases educating the participants on how- to sandbag (Higgins et al., 2017; Schatz & Glatts, 2013), the large differences of identified performance invalidity between EVI’s and PVT’s should not be ignored.

The discrepancies between rates of identified performance invalidity between EVIs and PVTs in this sample raise several possibilities. First, it provides support for the developing idea that performance validity is on a continuum where valid, optimal performance is on one end, intentional underperformance and invalid data is on the other, and less-than-valid (and potentially variable, disengaged, distracted, or apathetic) performance is in the middle. These results may also suggest that less-than-valid performance looks different depending on the motivation (or maybe lack thereof), with malingering (intentional poor performance with external incentive) and sandbagging (intentional poor performance on baseline in order to look recovered more quickly after concussion) demonstrating more egregious differences on test scores including validity measures and representing a more extreme end of the continuum.

These findings would also support the idea that using multiple validity checks across an evaluation is important and failed validity measures do not necessarily indicate intentional underperformance, but can only speak to the validity of the related data which may be influenced by a myriad of factors. ImPACT composites were significantly different based on the number of validity measures that an athlete was flagged on, but differences were not seen until the athlete flagged on multiple validity measures (2+ for Visual-Motor Speed composite, Verbal Memory composite proxy, Visual Memory composite proxy, and 3+ for Reaction Time composite). Additionally, this study provides a new and more methodologically sound framework for examining the validity of baseline data in athletes compared to the current commonly examined experimental malingering paradigm.

Also similar to previous research (Cook et al., 2017; Gaudet & Weyandt, 2017; Maietta et al., 2021; Maietta et al., 2023; Manderino & Gunstad, 2018; Manderino et al., 2019; Nelson et al., 2016; Schatz et al., 2014; Szabo et al., 2013; Tsushima et al., 2019), self-reported ADHD and/or LD diagnoses increased the number of validity measures flagged. These athletes also reported more symptoms (and less sleep), and demonstrated lower scores on cognitive composites. It should be noted that 19% of our sample self-reported a diagnosis of ADHD and/or LD, which is higher than population estimates. It is possible that our subgroup of athletes with ADHD and/or LD is not necessarily representative of the general population prevalence of the disorder in athletes. It is also possible that self-reported diagnoses were inaccurate and inflated the presence of ADHD and/or LD in our sample. The ongoing challenge in studying performance validity in athletes in self-reported ADHD and/or LD is differentiating invalid data from expected variability in cognitive performance due to diagnosis (e.g., variable and inefficient attention or reading difficulties). Future research using clinically confirmed diagnoses is certainly needed in this regard.

## Limitations

There were several limitations to this study. First ImPACT was completed in a group setting while the stand-alone PVTs were administered in an individual setting. Research has shown that administration setting can make a difference in ImPACT composite scores (Lichtenstein et al., 2014; Moser et al., 2011), so this may play a role as athletes may have put forth more effort and have few environmental distractions on PVT’s as they were one-on-one with an administrator.

Reliance on self-reported neurodevelopmental disorder and hours of sleep the night before testing are also limitations as compared to more objective data such as clinician confirmed diagnosis. The vast majority of the literature on athletes with ADHD and/or LD relies on self-reported diagnosis as compared to expert clinical diagnosis (Maietta et al., 2023). This is not only a limitation in our study, but also in the extant literature in this area. Future studies should strive to rely on confirmed clinical diagnoses when evaluating athletes with neurodevelopmental conditions such as ADHD or LD. Finally, our sample of athletes with self-reported ADHD and/or LD was relatively small, and the subset of the sample that also failed multiple validity measures was even smaller. This may limit the generalizability of these findings.

In addition, there may be differences in performance on ImPACT across sports (for example, contact and collision sport athletes being more likely to have completed multiple baseline and possibility post-injury tests and/or non-contact sport athletes having limited reason to sandbag, but also limited reason to want to perform well given the low likelihood of experiencing a sport-related concussion). Given that this sample of athletes was from 14 different sports we were unable to discern the effect sport may have on results.

## Future Directions

Two primary areas for future study were identified by this research. The first is continuing to study validity along a continuum, rather than using the typical dichotomous malingering/sandbagging paradigm. The validity of baseline data is paramount as it is often used as a benchmark for return-to-sport, and therefore head injury exposure. Also, baseline data is often interpreted by individuals without training in psychometrics or test interpretation, so sensitive and unambiguous markers for valid or invalid data need to be clearly operationalized and disseminated. An EVI-based model for evaluating performance validity has particular applicability to concussion baseline testing, as it is very often not feasible to add an additional stand-alone PVT. Finally, based on this data, as well as previous research, EVIs seem to be more sensitive to performance invalidity in concussion baseline testing than stand-alone PVTs.

The second area for ongoing research is continuing to study the most sensitive and specific ways to measure performance validity in neurodiverse athletes. The challenge in measuring performance validity using an EVI model in this population lies in an EVI model being based on lower-than-expected or more-variable-than-expected scores. Unfortunately, neurodiverse athletes may produce both lower and more variable scores than their neurotypical peers due to their diagnosis and so being able to pull apart data that is invalidity from data that is valid but negatively impacted by a developmental diagnosis is of utmost importance if baseline testing is going to have utility in this subset of athletes.

Finally, this article adds another data point in the growing body of literature that questions the utility (both in terms of reliability and return-on-investment) of baseline cognitive testing for managing sport concussion return-to-play decisions (Kirkwood et al., 2009). Baseline testing does not function any better than normative data in identifying post-injury change in many cases (Echemendia et al., 2012) and may actually increase risk of misclassification in some athlete groups (Schatz & Robertshaw, 2014), but is typically a very time- and resource-intense process (Iverson & Schatz, 2015). If as large of a percentage of baseline data has questionable validity as has been suggested by this and other research (Higgins et al., 2018), the baseline test model quickly becomes untenable. Alternatively, the answer may not be to do away with baseline testing, but to change the paradigms that we use to evaluate baseline functioning and performance validity. This could include conducting multiple trials of the same task and evaluating consistency across trials. Another possibility is creating less time-consuming (i.e., shorter, but more sensitive, and able to be administered in larger groups or remotely) but more frequently administered baseline evaluations, as research has demonstrated that baseline scores are increasingly reliable when averaged over a series of administrations (Bruce et al., 2016). A final possibility is generating new baseline cognitive measures that have more ecological validity or utility across disciplines (academics, sport performance psychology, and concussion management, etc).

